# Anti-severe acute respiratory syndrome-related coronavirus 2 (SARS-CoV-2) potency of Mefloquine as an entry inhibitor in vitro

**DOI:** 10.1101/2020.11.19.389726

**Authors:** Kaho Shionoya, Masako Yamasaki, Shoya Iwanami, Yusuke Ito, Shuetsu Fukushi, Hirofumi Ohashi, Wakana Saso, Tomohiro Tanaka, Shin Aoki, Kouji Kuramochi, Shingo Iwami, Yoshimasa Takahashi, Tadaki Suzuki, Masamichi Muramatsu, Makoto Takeda, Takaji Wakita, Koichi Watashi

## Abstract

Coronavirus disease 2019 (COVID-19) has caused serious public health, social, and economic damage worldwide and effective drugs that prevent or cure COVID-19 are urgently needed. Approved drugs including Hydroxychloroquine, Remdesivir or Interferon were reported to inhibit the infection or propagation of severe acute respiratory syndrome-related coronavirus 2 (SARS-CoV-2), however, their clinical efficacies have not yet been well demonstrated. To identify drugs with higher antiviral potency, we screened approved anti-parasitic/anti-protozoal drugs and identified an anti-malarial drug, Mefloquine, which showed the highest anti-SARS-CoV-2 activity among the tested compounds. Mefloquine showed higher anti-SARS-CoV-2 activity than Hydroxychloroquine in VeroE6/TMPRSS2 and Calu-3 cells, with IC_50_ = 1.28 μM, IC_90_ = 2.31 μM, and IC_99_ = 4.39 μM in VeroE6/TMPRSS2 cells. Mefloquine inhibited viral entry after viral attachment to the target cell. Combined treatment with Mefloquine and Nelfinavir, a replication inhibitor, showed synergistic antiviral activity. Our mathematical modeling based on the drug concentration in the lung predicted that Mefloquine administration at a standard treatment dosage could decline viral dynamics in patients, reduce cumulative viral load to 7% and shorten the time until virus elimination by 6.1 days. These data cumulatively underscore Mefloquine as an anti-SARS-CoV-2 entry inhibitor.

## 1. Introduction

Coronavirus disease 2019 (COVID-19), caused by infection of severe acute respiratory syndrome-related coronavirus 2 (SARS-CoV-2), has spread into a worldwide since it was first reported in Wuhan, China in December 2019, and caused severe damage to public health, the economy, and society in many countries and areas. Several therapeutic drug candidates, including Remdesivir (RDV), Hydroxychloroquine (HCQ), Lopinavir and Interferon, have been undergone clinical trials with drug-repurposing approaches (Touret et al., 2020), of which treatment efficacies have yet been fully demonstrated. New drug choices for both therapeutic and prophylactic use against COVID-19 are urgent needs.

Chloroquine and its derivative, HCQ, are used clinically as anti-malarial drugs (Sinha et al., 2014). These drugs (particularly the less toxic HCQ) were expected to be COVID-19 drug candidates from the early days of the COVID-19 pandemic (Cortegiani et al., 2020), given their anti-SARS-CoV-2 activity *in vitro* and the ability to reduce pathogenesis caused by the related coronaviruses, SARS-CoV and human coronavirus OC43 *in vivo* (Keyaerts et al., 2009; Weston et al., 2020; Wang et al., 2020; Liu et al., 2020). However, despite over 30 randomized controlled trials or observational studies in different countries, no consensus demonstrates a sufficient anti-COVID-19 effect of these drugs (Geleris et al., 2020; Rosenberg et al., 2020; Tang et al., 2020; Yu et al., 2020a). Therefore, the FDA revoked the emergency use of chloroquine and HCQ for COVID-19 treatment in June 2020. The discrepancy between *in vitro* and *in vivo* experimental data and the clinical outcomes reported to date is not well understood. Possibilities include differences in drug sensitivities among cell types used in experiments (see *4. Discussion*) and the insufficient potential of anti-SARS-CoV-2 activity of these drugs: The concentrations of HCQ required for 50% and 90% virus reduction (IC_50_, IC_90_), determined *in vitro* (i.e., several μM), is higher than an achievable in plasma value in clinical settings (1-2 μM at the maximum) (McLachlan et al., 1993; Touret et al., 2020; Liu et al., 2020; Hattori et al., 2020) (see *4. Discussion*). Thus, identifying another drug with a higher antiviral potential at the maximum drug concentration based on clinical data is a probable approach to improving the treatment efficacy. In this study, from a cell-based functional screening of FDA/EMA/PMDA-approved anti-parasitic/anti-protozoal drugs, we identified Mefloquine (MFQ), a derivative of HCQ originally used for anti-malarial therapy and prophylaxis (Sinha et al., 2014), that has a higher anti-SARS-CoV-2 activity than HCQ in both TMPRSS2-overexpressed VeroE6 cells and human lung-derived Calu-3 cells. MFQ inhibited viral entry process after attachment of the virus to the cell. Importantly, our mathematical modeling predicted that MFQ administration (1,000 mg, once per day) could decline viral dynamics in patients to significantly reducing the cumulative viral load and shortening the period until virus elimination in clinical concentration ranges. Our data provide foundational evidence that proposes MFQ as an alternative drug for anti-COVID-19 treatment.

## 2. Materials and Methods

Information for Materials and Methods are described in ***Supplementary Information***.

## 3. Results

### 3.1. Identification of Mefloquine as a potential inhibitor against SARS-CoV-2 infection

In this study, we mainly used VeroE6/TMPRSS2 cells, which is established by overexpressing transmembrane serine protease 2 (TMPRSS2) in VeroE6 cells (Nao et al., 2019; Matsuyama et al., 2020), and human lung epithelial-derived Calu-3 cells in a part of experiments, as SARS-CoV-2 infection models. First, we examined the dose dependency of HCQ for antiviral activity by a cytopathic effect (CPE) assay: VeroE6/TMPRSS2 cells were inoculated with SARS-CoV-2 at an MOI of 0.001 for 1 h, washed to remove unbound virus, and incubated for an additional 48 h (Fig. 1A). SARS-CoV-2 propagation in the cells exhibited an intensive cytopathic effect (Fig. 1B, panel b), as reported (Matsuyama et al., 2020). HCQ protected cells from SARS-CoV-2-induced cytopathology completely at the concentration of 32 μM, remarkably but not completely at 16 μM, and very little at 8 μM (Fig.1B, panels c-e).

**Figure. 1.**
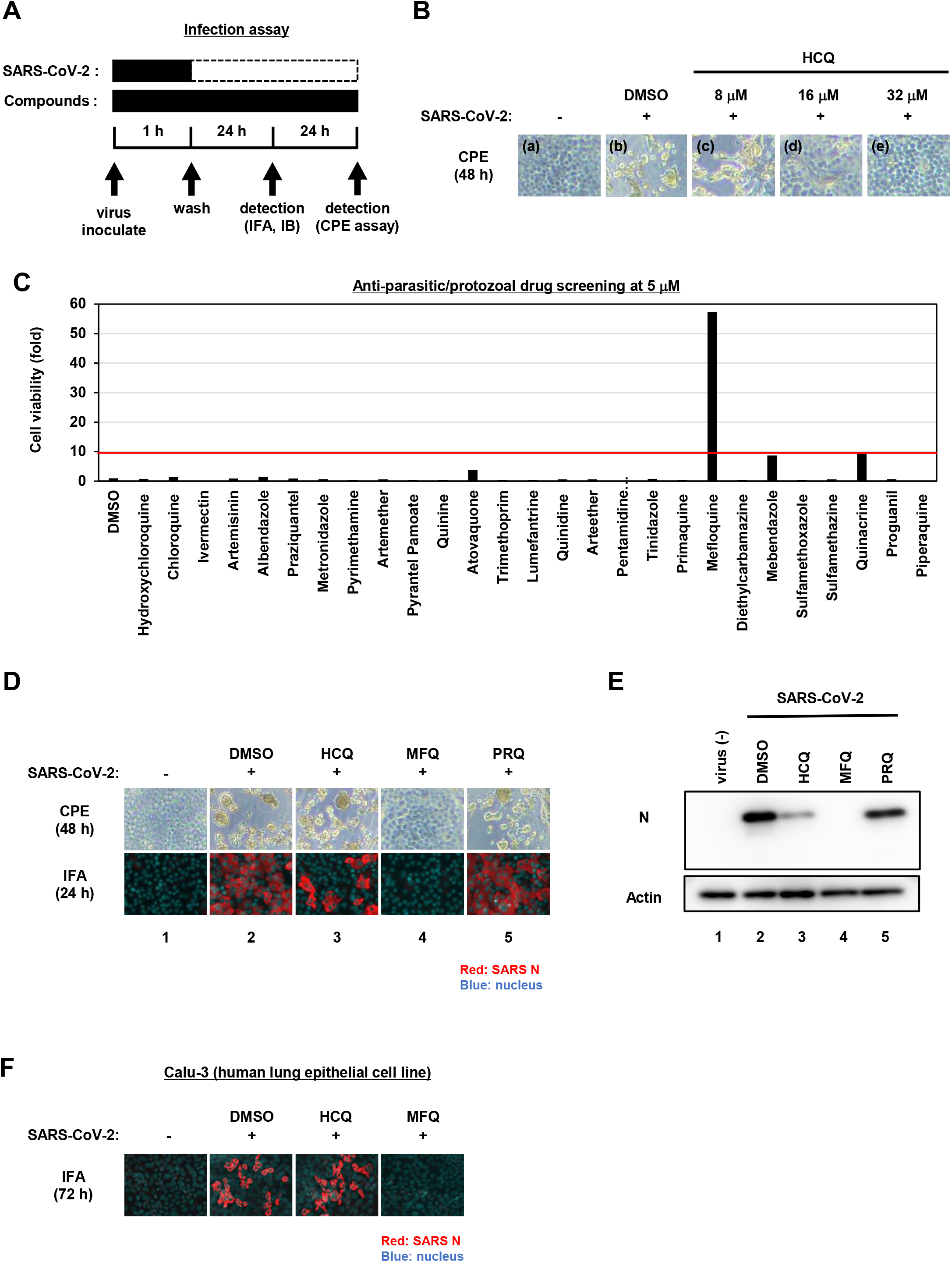
Mefloquine (MFQ) inhibits Severe Acute Respiratory Syndrome-related coronavirus 2 (SARS-CoV-2) propagation. **(A)** Schematic representation of the SARS-CoV-2 infection assay. VeroE6/TMPRSS2 cells were inoculated with SARS-CoV-2 (Wk-521 strain) at an MOI of 0.001 for 1 h. After removing the unbound virus, cells were cultured for 24 h to detect virus-encoding N protein by immunofluorescence assay (IFA) and immunoblot (IB) or to detect viral RNA in the culture supernatant by RT-qPCR, or for 48 h to observe virus-induced cytopathic effect (CPE). Compounds were treated given throughout the assay. **(B)** Dose dependency of Hydroxychloroquine (HCQ) on CPE suppression. VeroE6/TMPRSS2 cells were inoculated with the virus for 1 h. Removing the unbound virus, cells were cultured with a medium containing the indicated compounds for 48 h. CPE was observed by microscopy. **(C)** Screening of anti-parasitic/protozoal drugs in the cell-based infection assay. Compounds were administrated at 5 μM, at which hydroxychloroquine showed little effect on CPE. The viability of infected cells was quantified via a high content imaging analyzer by setting the value for the sample treated with DMSO solvent as 1. MFQ showed more than 57-fold higher cell viability than DMSO controls. **(D, E)** SARS-CoV-2-induced CPE and viral N protein expression upon compound treatments [DMSO at 0.08%; hydroxychloroquine (HCQ), mefloquine (MFQ), and primaquine (PRQ) at 8 μM]. Red and blue signals of merged images indicate viral N protein and nucleus, respectively (D, lower). Viral N protein and actin, an internal control, were detected by immunoblot (E). **(F)** The anti-SARS-CoV-2 activity of the indicated compounds in Calu-3 cells, a human lung epithelial cell-derived line.

Aiming to identify drugs with greater anti-SARS-CoV-2 potential than HCQ, we employed 5 μM for drug screening, a concentration at which HCQ had no CPE suppression. As a drug library, we used approved anti-parasitic/anti-protozoal drugs for following two reasons; 1) In addition to Chloroquine and HCQ, some drugs such as Ivermectin, Atovaquone and quinoline derivatives were reported to demonstrate antiviral activities against other RNA viruses (Cifuentes Kottkamp et al., 2019; DeWald et al., 2019; Mastrangelo et al., 2012; Al-Bari, 2015). 2) Anti-parasitic/anti-protozoal agents generally reach high concentrations (i.e., over μM ranges) in the plasma in clinical settings (Sinha et al., 2014). We thus screened 27 FDA/EMA/PMDA-approved (or approved in the past) anti-parasitic/anti-protozoal drugs at 5 μM by the CPE assay (Fig. 1A, *Supplementary Materials and Methods*). By following the scheme shown in Fig. 1A, cells at 48 h post-inoculation were fixed, stained with DAPI, and counted to quantify surviving cell numbers. The graph in Fig. 1C shows survival cell numbers relative to that of DMSO-treated cells as a control, and survival cell number relative to that of non-infected cells are shown in Fig. S1. In this screening, HCQ, Chloroquine and Ivermectin had little effect, while MFQ remarkably protected cells from the virus-induced CPE, with a more than 57-fold increase in surviving cells over those of the vehicle control (Fig. 1C).

We next compared the antiviral activities of MFQ with that of HCQ and an additional Chloroquine derivative, Primaquine (PRQ), as a reference. Cytopathogenicities at 48 h and the viral N protein expression at 24 h after virus inoculation (a time before showing CPE) were examined during treatment with each compound at 8 μM (Fig. 1D, E): MFQ completely protected cells from viral propagation-induced CPE and reduced the production of viral protein (lane 4), whereas HCQ weakly exerted an antiviral effect (lane 3), and PRQ had little antiviral effect (lane 5). To examine whether the observed antiviral effects depend on cell types or are generally reproduced beyond cell types, we used a human lung epithelial cell line, Calu-3, and found the robust antiviral activity of MFQ against SARS-CoV-2, in contrast to much lower HCQ activity (Fig. 1F, *Supplementary Materials and Methods*). Therefore, we focused on MFQ as a potential anti-SARS-CoV-2 drug in subsequent analyses.

### 3.2. Antiviral profile of Mefloquine and other quinoline derivatives

To profile the anti-SARS-CoV-2 activity of compounds, we quantified viral RNA released into the culture supernatant in addition to cell viability at 24 h after virus inoculation upon treatment at varying concentrations (0.5, 1, 2, 4, 8 and 16 μM) of HCQ, PRQ, MFQ, and other related compounds, Quinine and Quinidine, that possess a quinoline ring (Fig. 2A–C). The 90% and 99% maximal inhibitory concentrations (IC_90_ and IC_99_) and 50% maximal cytotoxic concentrations (CC_50_) are shown. All the compounds had no remarkably cytotoxicity at any examined concentration (Fig. 2C). HCQ and MFQ demonstrated antiviral activities in a dose-dependent manner, with higher potency for MFQ than HCQ (Fig. 2B). By contrast, PRQ showed marginal antiviral effects at all concentrations examined, suggesting that the hydroxyl and amino groups in the side chain of MFQ and/or that the position of the side chain on the quinoline ring are important for the anti-SARS-CoV-2 activity. The octanol-water partition coefficient (log P) values of MFQ, HCQ, Quinine, Quinidine and PRQ were calculated to be 4.34, 2.87, 2.48, 2.4, and 1.47, respectively (Ghose and Crippen, 1987), which imply that the higher hydrophobicity of MFQ, possibly due to the two trifluoromethyl groups, may be related to its high antiviral activity.

**Figure. 2.**
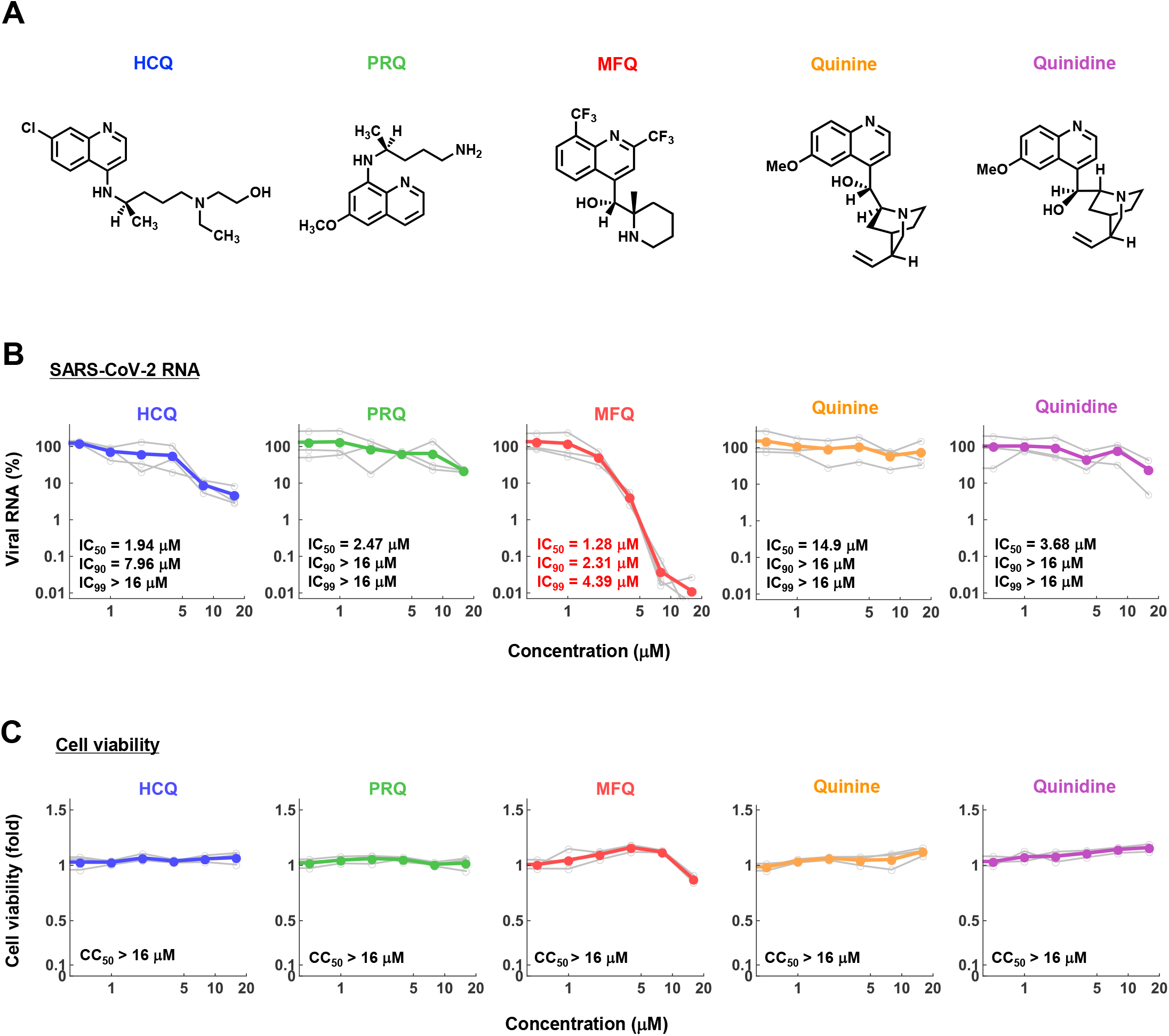
The anti-SARS-CoV-2 activity of MFQ and its derivatives. **(A)** Chemical structures of MFQ and its derivatives. **(B)** Extracellular SARS-CoV-2 RNA was quantified upon treatment with HCQ, MFQ and related compounds PRQ, Quinine and Quinidine at varying concentrations. Calculated inhibitory concentrations of 50%, 90% and 99% maximum (IC_50_, IC_90_ and IC_99_) for each compound are as indicated. **(C)** Cell viability was measured by MTT assay with the calculated 50% maximal cytotoxic concentration (CC_50_).

### 3.3. Mefloquine inhibits the SARS-CoV-2 entry process after virus-cell attachment

SARS-CoV-2 attaches to target cells by the binding of viral Spike protein to its receptor, angiotensin-converting enzyme 2 (ACE2). It is then subjected to Spike cleavage by host proteases, either TMPRSS2 on the plasma membrane or cathepsins in the endosomes, followed by the membrane fusion and the sorting to the site of replication (entry phase). Viral RNA then replicates and assembles with viral structural proteins to produce progeny virus (replication phase) (Fig. 3A) (Hoffmann et al., 2020; Lebeau et al., 2020).

**Figure. 3.**
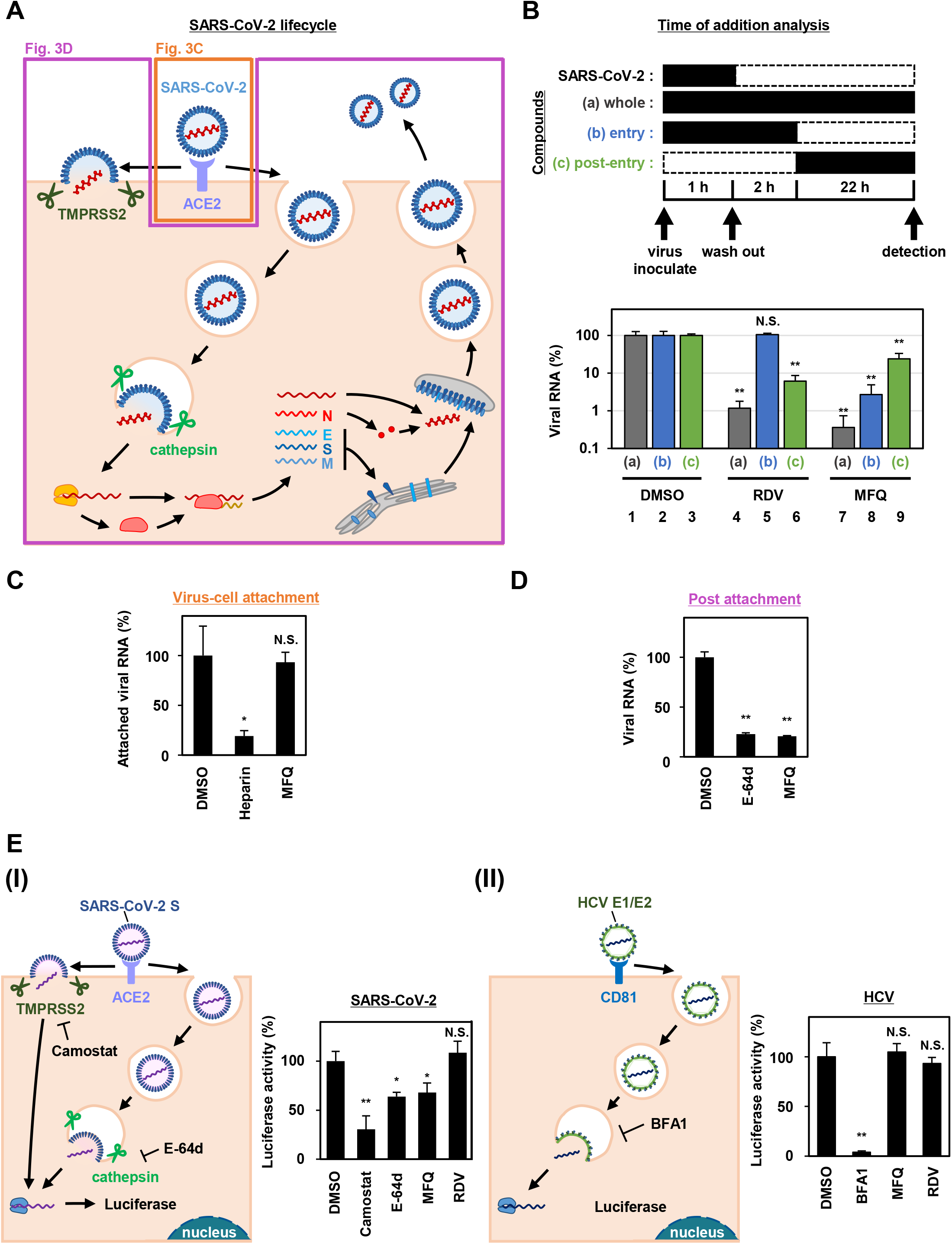
MFQ inhibits the SARS-CoV-2 entry process. **(A)** SARS-CoV-2 life cycle. SARS-CoV-2 infection is initiated with virus attachment to the host cells that involves the cellular receptor, angiotensin converting enzyme 2 (ACE2), followed by the cleavage of viral Spike (S) proteins by either transmembrane serine protease (TMPRSS2) on the plasma membrane or cathepsins in the endosome/lysosome that induces fusion of viral and host membranes. Viral RNA is translated, processed and replicated to be assembled into progeny virus with viral structural proteins and released extracellularly. **(B)** Scheme of the time of addition analysis. Compounds were treated at three different times: **(a) whole**: throughout the assay for 25 h, **(b) entry**: for the initial 3 h to evaluate the effect on the viral entry process and **(c) post-entry**: for the last 22 h to evaluate the effect on viral replication/re-infection. Viral RNA levels in the culture supernatant are shown in the graph by setting that upon DMSO treatment as 100%. **(C)** Virus-cell attachment assay. VeroE6/TMPRSS2 cells were exposed to virus at an MOI of 0.001 at 4°C for 5 min with 50 μM MFQ or 100 U/mL Heparin, a SARS-CoV-2 attachment inhibitor used as a positive control. After washing the unbound virus, cell surface-attached virus was extracted and quantified by real-time RT-PCR. **(D)** Post-attachment assay. For evaluating the activity after virus attachment, from membrane fusion to virus secretion, VeroE6/TMPRSS2 cells preincubated with the virus at an MOI of 1.5 at 4°C for 1 h to allow virus attachment were treated with compounds for 6 h at 37°C. Extracellular viral RNA was quantified by RT-qPCR. E-64d, a cysteine protease inhibitor, was used as a positive control. **(E)** Pseudovirus assays carrying the SARS-CoV-2 Spike or hepatitis C virus (HCV) E1E2 envelope. In the SARS-CoV-2 pseudovirus assay, Camostat and E-64d were used as positive controls for inhibiting TMPRSS2 and cysteine protease, respectively (E, left). Bafilomycin A1 (BFA1), which reported to inhibit HCV entry, was used as a positive control for HCV pseudovirus assay (E, right).

We next addressed which step in the viral life cycle MFQ inhibits by a series of assays. The time-of-addition analysis, in which compounds are treated at different times, is used to evaluate the phase of viral entry and replication separately (Wang et al., 2020). As previously reported (Wang et al., 2020), compounds were treated at three different time points (Fig. 3B, *Supplementary Materials and Methods*), either throughout the assay (a; whole life cycle, 1 h during virus inoculation + 24 h after inoculation), for the initial 3 h (b; entry phase, 1 h during virus inoculation + 2 h after inoculation), or for the last 22 h (c; post-entry phase, including replication). In this analysis, RDV, a reported replication inhibitor (Wang et al., 2020), had no inhibitory effect when applied during the initial 3 h (Fig. 3B, lane 5), but it decreased viral RNA in the post-entry phase (Fig. 3B, lane 6). By contrast, MFQ remarkably reduced viral RNA levels to under 3% when applied at the entry phase (Fig. 3B, lane 8), but showed much lower antiviral activity (to 24%) when treated after the first round of viral entry (Fig. 3B, lane 9). The viral RNA reduction by MFQ in lane 9 was likely to the inhibition of second round of infection and thereafter of the produced virus, which occurred during the 22 h. These data suggest that MFQ inhibits the entry process of SARS-CoV-2.

We then evaluated the virus-cell attachment in the presence or absence of MFQ by incubating cells with the virus at 4°C to allow viral attachment to the cell surface but not the following processes. After washing the unattached virus and compounds, we extracted and quantified the viral RNA on the cell surface. SARS-CoV-2 RNA from virus attached the surface of the cell was drastically reduced in the presence of heparin, an entry inhibitor for SARS-CoV-2, used as a positive control (Tandon et al., 2020; Tree et al., 2020), while that was not affected by MFQ treatment (Fig. 3C). However, MFQ inhibited the post-attachment phase, ranging from the membrane fusion to virus production (Fig. 3D): Virus-attached cells were prepared by incubation with a large amount of virus (MOI of 1.5, more than 1,000-fold higher than used in other normal infection assay) at 4°C for 1 h followed by washing. The cells were transferred to 37°C for 6 h in the presence or absence of compounds to induce membrane fusion and subsequent steps up to virus secretion, and viral RNA in the supernatant was quantified. MFQ clearly reduced the viral RNA levels to almost the same as those when treatment with E-64d, a lysosomal/cytosolic cysteine protease inhibitor reported to inhibited SARS-CoV-2 entry (Hoffmann et al., 2020; Hu et al., 2020) (Fig. 3D).

We further examined the virus entry using a pseudovirus carrying the Spike protein derived from SARS-CoV-2 or the envelope proteins of hepatitis C virus (HCV), another RNA virus unrelated to coronavirus (Fig. 3E, *Supplementary Materials and Methods*). These pseudoviruses can evaluate the entry mediated by these Spike or envelope proteins (Hoffmann et al., 2020; Bartosch et al., 2003). The pseudovirus assay showed that SARS-CoV-2 Spike-dependent viral entry was significantly inhibited by the TMPRSS2 inhibitor Camostat, and by MFQ to similar levels to those of E-64d (Fig. 3E, left). However, the assay sensitivity itself was relatively lower than the SARS-CoV-2 infection assay. Meanwhile, HCV envelope-mediated entry was not affected by MFQ, in contrast to the reduced entry caused by bafilomycin A1, a reported HCV entry inhibitor (Fig. 3E, right). These results cumulatively suggest that MFQ inhibited the post-attachment SARS-CoV-2 Spike-dependent entry process.

### 3.4. Synergistic antiviral activity of combined treatment of Mefloquine with Nelfinavir

Combination treatment with multiple agents with different modes of action is a strategy to improve the outcome of antiviral treatments, including those against human immunodeficiency virus (HIV) and HCV (Koizumi et al., 2017; Shen et al., 2008). We, therefore, examined the combination of MFQ and a representative anti-SARS-CoV replication inhibitor, Nelfinavir (NFV) (Yamamoto et al., 2004). NFV has been suggested to inhibit SARS-CoV-2 replication thorough binding with the SARS-CoV-2 main protease by docking simulation (Ohashi et al., 2020). Following the experimental scheme in Fig. 1A, we treated cells with paired compounds at varying concentrations for 24 h and quantified viral RNA in the cultured supernatant by real-time RT-PCR in addition to cell viability by a high content image analyzer (*Supplementary Materials and Methods*). Viral RNA levels were reduced by a single treatment of either MFQ or NFV in a dose-dependent manner, and these was further reduced by combination treatment without any cytotoxicity (Fig. 4A). Bliss independence-based synergy plot showed a synergistic antiviral effect in wide concentration ranges, especially at higher doses (Fig. 4B, orange indicates synergistic effect).

**Figure. 4.**
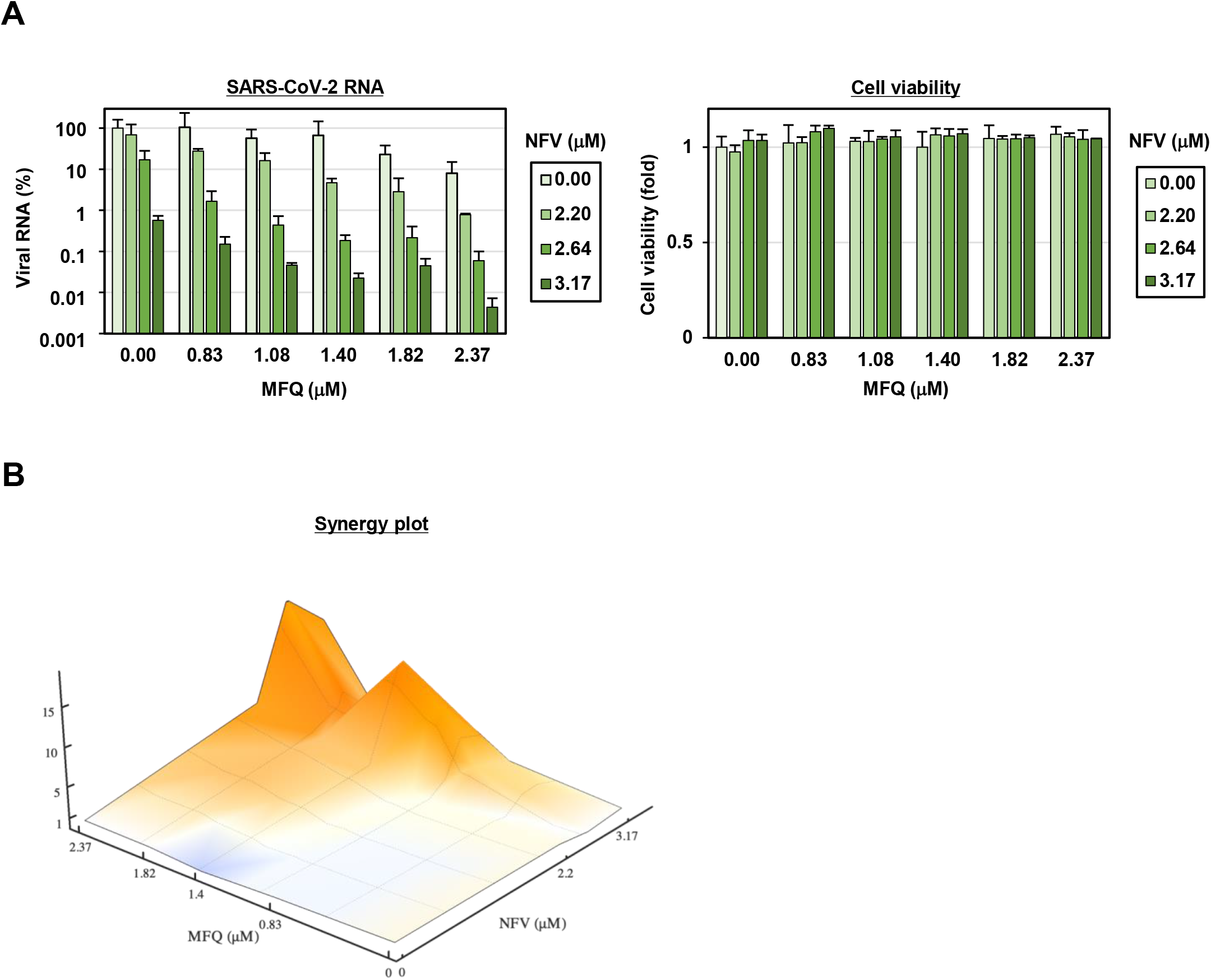
MFQ shows synergistic anti-SARS-CoV2 activity with replication inhibitor NFV. **(A)** Viral RNAs in the culture supernatant at 24 h after co-treatment with MFQ and NFV were quantified by real-time RT-PCR. Relative values are shown of viral RNA or cell viability to those treated with DMSO control. Cell viability was simultaneously measured with a high content image analyzer. [MFQ at 0, 0.83, 1.08, 1.40, 1.82 and 2.37 μM (1.3-fold-dilution); NFV at 0, 2.20, 2.64 and 3.17 μM (1.2-fold-dilution)]. **(B)** The three-dimensional interaction landscapes of NFV and MFQ were evaluated with the Bliss independence model. Orange, white and dark-blue colors on the contour plot indicate synergy, additive and antagonism, respectively.

### 3.5. Mathematical prediction of the Mefloquine treatment in clinical settings

Clinical pharmacokinetics data for MFQ, including the maximum drug concentration (C_max_) in the plasma, half-life, area under the curve for drug concentration, and the distribution to the lung, are reported (Desjardins et al., 1979; Karbwang and White, 1990; Jones et al., 1994). Mathematical modeling combined with pharmacokinetics, pharmacodynamics, and the viral dynamics model described in **Materials and Methods** (Ohashi et al., 2020) predicted the dynamics of viral load after MFQ administration (1,000 mg, once) in patients (Fig. 5A, red) and the corresponding time-dependent antiviral activity of MFQ (Fig. 5B). The high antiviral potential and the long half-life of MFQ (more than 400 h) (Desjardins et al., 1979; Karbwang and White, 1990) were predicted to exert a continuous antiviral effect and a resulting decline of viral load (Fig. 5A). Cumulative viral load, which is the area under the curve for the viral load over the time course, was calculated to be reduced by 6.98% (Fig. 5C). The time until the viral load declines beneath the detectable level is 15.2 days without treatment, but it was calculated to be shortened to 9.10 days after MFQ treatment (Fig. 5D). These analyses predict the effectiveness of MFQ to reduce the viral load at clinical drug concentrations.

**Figure. 5.**
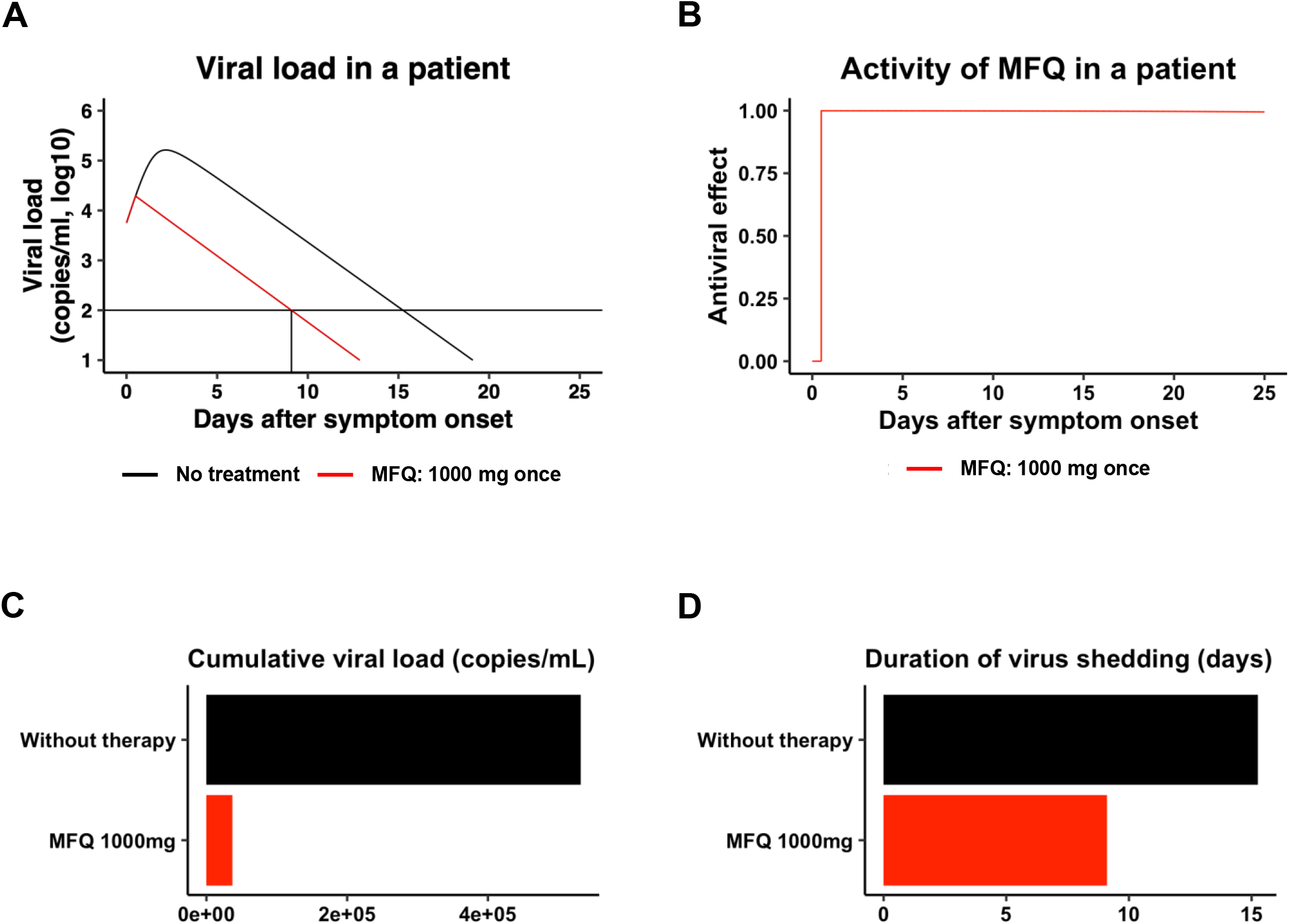
Prediction of the impact of MFQ treatment on SARS-CoV-2 dynamics in clinical settings. **(A, B)** The predicted viral load dynamics without (A, black) or upon MFQ administration (1,000mg oral, once per day) (A, red) and the time-dependent antiviral activity of MFQ (B) predicted by pharmacokinetics/pharmacodynamics/viral-dynamics (PK/PD/VD) models. **(C, D)** The cumulative viral load calculated as the area under the curve in **(A)** and the duration of virus shedding (days) [time from symptom onset to the day achieving a viral load under the detection limit (black horizontal line) in **(A)**] were evaluated for nontreatment (black) or MFQ treatment (red).

## 4. Discussion

Given the *in vitro* anti-SARS-CoV-2 activity and the *in vivo* effect on the related coronaviruses (Ko et al., 2020; Weston et al., 2020; Wang et al., 2020; Liu et al., 2020), Chloroquine and HCQ have been expected to be effective as anti-COVID-19 drugs. However, accumulative data have not provided sufficient evidence supporting a preferable clinical outcome (Funnell et al., 2020). The IC_50_, IC_90_ and IC_99_ for HCQ calculated in this study were 1.94, 7.96 and 37.2 μM, respectively, consistent with the IC_50_ values at μM ranges examined in other studies (Liu et al., 2020; Touret et al., 2020; Gendrot et al., 2020; Hattori et al., 2020). Pharmacokinetics analyses in healthy volunteers receiving oral administration of 200 mg HCQ demonstrated a C_max_ in the blood of 0.49-0.55 μM (McLachlan et al., 1993), lower than the concentration ranges having significant anti-SARS-CoV-2 activity. These data led us to identify a drug possessing a greater anti-SARS-CoV-2 potential.

SARS-CoV-2 entry requires the initial binding of the viral Spike protein to its cell surface receptor ACE2, then Spike cleavage by either of the two independent host proteases, endosomal pH-dependent cathepsin or plasma membrane pH-independent TMPRSS2 (Hoffmann et al., 2020) (Fig. 3A). Recently, it has been reported that the sensitivity to viral entry inhibitors such as Chloroquine, HCQ and a TMPRSS2 inhibitor Camostat depends on cell types, so that recommended not to rely only on widely used Vero cell line, but to use rather TMPRSS2-complemented Vero cells, Calu-3 cells or presumably primary respiratory/lung cell culture in an air-liquid interface system or organoids as a more physiologically relevant model for airway epithelial cells (Hoffmann et al., 2020; Suzuki et al., 2020). Due to the poor availability of primary cells, we employed VeroE6/TMPRSS2 and Calu-3 cells in this study, and discovered that MFQ inhibited the viral entry more potently than HCQ in these TMPRSS2-expressing cells. Importantly, standard MFQ treatment given to healthy volunteers achieved a plasma C_max_ of 4.58 μM with a long half-life (more than 400 h) (Karbwang and White, 1990), which is within concentration ranges exerting significant anti-SARS-CoV-2 activity *in vitro*. Moreover, it has been reported that the MFQ concentration in the lung was over 10-fold that of the blood in MFQ-treated human participants (Jones et al., 1994), expecting an even higher anti-SARS-CoV-2 effect of MFQ. Our mathematical model analysis (Fig. 5) quantified this prediction, demonstrating a clear reduction in both cumulative viral load in patients and the time for viral elimination.

The *in vitro* anti-SARS-CoV-2 activity of MFQ itself has been reported (Fan et al., 2020; Jeon et al., 2020; Gendrot et al., 2020; Weston et al., 2020), however, they only reported the anti-SARS-CoV-2 activity in a single cell line (Vero or VeroE6 cells) with a single readout (viral RNA or CPE) at only one experimental condition without mechanistic analysis. In the present study, in addition to the comparing the activity of MFQ with HCQ and other analogs side-by-side, we characterized the modes of action and combination treatments. Furthermore, we addressed the clinical antiviral efficacy of MFQ by mathematical prediction, a significant scientific novelty. Our time-of-addition, virus-cell attachment, post attachment and pseudovirus assays suggest that MFQ inhibits the SARS-CoV-2 entry phase after attachment, including the viral Spike cleavage/membrane fusion and the following translocation to the replication complex. Detailed analysis of the mode of action is the object of future studies.

A limitation of our study is the use of antiviral profile data in cell culture assays but without an *in vivo* infection model. To date, SARS-CoV-2 studies have used models including hACE2-transgenic mice, ferrets, cats, hamsters, nonhuman primates and mice infected with mouse-adapted SARS-CoV-2 (Bao et al., 2020; Jiang et al., 2020; Hassan et al., 2020; Sun et al., 2020; Winkler et al., 2020; Golden et al., 2020; Kim et al., 2020; Shi et al., 2020; Richard et al., 2020; Sia et al., 2020; Imai et al., 2020; Rogers et al., 2020; Rockx et al., 2020; Gao et al., 2020; Yu et al., 2020b; Gu et al., 2020). However, except for antibodies or vaccine candidates, there are very limited reports at present successfully confirming the reduction of SARS-CoV-2 viral load in these models by treatment with drug candidates (Park et al., 2020). At this time, however, proposing an additional treatment choice with significant antiviral evidences is urgently demanded to combat COVID-19. Interestingly, MFQ showed a synergistic effect combined with a replication inhibitor for SARS-associated coronavirus, NFV (Yamamoto et al., 2004; Ohashi et al., 2020) (Fig. 4). These data would prospect better clinical outcomes by combined drugs with different modes of action, as used with antiviral therapy against HIV and HCV (Koizumi et al., 2017; Shen et al., 2008). Given the inhibition of viral entry, MFQ is also expected for prophylactic use. Its long half-life of approximately 20 days is advantageous for achieving a long-lasting antiviral state by a single oral administration. Consequently, our analysis highlights the anti-SARS-CoV-2 potency of MFQ, of which efficacy is expected to be further evaluated in the future through *in vivo* or clinical testing.

## Acknowledgments

We thank Dr. Shutoku Matsuyama at Department of Virology III of National Institute of Infectious Diseases in Tokyo and Dr. Shinichi Saito at Faculty of Sciences Division I of Tokyo University of Science for their technical assistance and discussion. The retrovirus-based pseudoparticle system and human hepatoma cell line, HuH-7 cells were kindly provided by Dr. Francois-Loic Cosset at the University of Lyon and Dr. Francis Chisari at The Scripps Research Institute, respectively.

