## Supplementary Information for "Anti-severe acute respiratory syndrome-related coronavirus 2 (SARS-CoV-2) potency of Mefloquine as an entry inhibitor in vitro"

[a]Department of Virology II, National Institute of Infectious Diseases, Tokyo 162-8640, Japan, [b]Department of Applied Biological Science, Tokyo University of Science, Noda 278-8510, Japan, [c]Department of Biology, Faculty of Sciences, Kyushu University, Fukuoka 812-8581, Japan, [d]Department of Virology I, National Institute of Infectious Diseases, Tokyo 162-8640, Japan, [e]The Institute of Medical Science, The University of Tokyo, Tokyo 108-8639, Japan, [f]AIDS Research Center, National Institute of Infectious Diseases, Tokyo 162-8640, Japan, [g]Faculty of Pharmaceutical Sciences, Tokyo University of Science, [h]Research Institute for Science and Technology, Tokyo University of Science, [i]MIRAI, JST, Saitama 332-0012, Japan, [j]Institute for the Advanced Study of Human Biology (ASHBi), Kyoto University, Kyoto 606-8501, Japan, [k]NEXT-Ganken Program, Japanese Foundation for Cancer Research (JFCR), Tokyo 135-8550, Japan, [l]Science Groove Inc., Fukuoka 810-0041, Japan, [m]Department of Immunology, National Institute of Infectious Diseases, Tokyo 162-8640, Japan, [n]Department of Pathology, National Institute of Infectious Diseases, Tokyo 162-8640, Japan, [o]Department of Virology III, National Institute of Infectious Diseases, Tokyo 208-0011, Japan, [p]Institute for Frontier Life and Medical Sciences, Kyoto University, Kyoto 606-8507, Japan.

Supplementary Materials and Methods
Supplementary Note
Supplementary Figure S1
Supplementary Table S1, S2
Supplementary References

### Supplementary Materials and Methods

*Cell culture.*

VeroE6/TMPRSS2 cells, a VeroE6 cell clone overexpressing the transmembrane protease, serine 2 gene (TMPRSS2) from Japanese Collection of Research Bioresources (JCRB) cell bank (Nao et al., 2019; Matsuyama et al., 2020), were cultured in Dulbecco's modified Eagle's medium (D-MEM; Life Technologies) supplemented with 10% fetal bovine serum (FBS; Cell Culture Bioscience), 10 units/mL penicillin, 10 mg/mL streptomycin, 10 mM HEPES (pH 7.4) and 1 mg/mL G418 (Nacalai) at 37°C in 5% $CO_2$. During the infection assay, G418 was removed and 10% FBS was replaced with 2% FBS. Calu-3 cells, a human lung epithelial cell line, were cultured in the above medium without G418 through the assay.

*Reagents.*

All the reagents were purchased from Selleck, Cayman Chemical and Tokyo Chemical Industry (TCI).

*Infection assay.*

SARS-CoV-2 was handled in a biosafety level 3 (BSL3) facility. We used the SARS-CoV-2 Wk-521 strain, a clinical isolate from a COVID-19 patient, that was propagated in VeroE6/TMPRSS2 cells and amplified (Matsuyama et al., 2020). Virus infectious titer ($TCID_{50}$/mL) was measured by observing the cytopathic effect of cells inoculated with 10-fold serial dilution of the virus (Matsuyama et al., 2020). For the infection assay using VeroE6/TMPRSS2 cells, SARS-CoV-2 was inoculated at a multiplicity of infection (MOI) of 0.001 for 1 h, and the unbound virus was removed by washing (Fig. 1B-E, 2B, 3B-C, 5A). Cells were cultured for 24 h to measure extracellular viral RNA or to detect viral N protein, or for 48 h to observe virus-induced cytopathic effect (CPE). Compounds were added during virus inoculation (1 h) and after inoculation (24 or 48 h), except the time-of-addition assay (Fig. 3B) and the assay evaluating the post-attachment phase from membrane fusion to virus secretion (Fig. 3D).

The Calu-3 cell infection assay was performed by virus inoculation at an MOI of 0.1 for 3 h and incubation for an additional 72 h to detect viral N protein (Fig. 1F).

*Compound screening.*

We screened 27 approved anti-parasitic and anti-protozoal drugs (Selleck). VeroE6/TMPRSS2 cells were treated with 5 μM of each drug for 1 h during virus inoculation at an MOI of 0.001. After removing the unbound virus, the cells were incubated with the drugs for an additional 48 h and were recovered, fixed in 4% paraformaldehyde and stained with 0.02% 4′, 6-diamidino-2-phenylindole (DAPI). The number of surviving cells was quantified with a high content imaging analyzer. The survival cell numbers treated with each drug are presented as a fold value relative to the cells treated with DMSO solvent (Fig. 1C). Drugs that protected cells from virus-induced CPE to more than 10-fold of the infected cells treated with DMSO was selected as hits.

*Immunofluorescence and Immunoblot analysis.*

Viral encoded N protein expression was detected using a rabbit anti-SARS-CoV N antibody (Mizutani et al., 2004) as a primary antibody with anti-rabbit AlexaFlour 568 or anti-rabbit IgG-HRP (ThermoFisher) as a secondary antibody by indirect immunofluorescence or immunoblot analysis (Fig. 1D-F) as previously reported (Ohashi et al., 2018). Anti-actin (Sigma) was used as an internal control for the immunoblot analysis. For immunofluorescence, nuclei were stained with DAPI (blue).

*Quantification of viral RNA.*

Viral RNA was extracted with QIAamp Viral RNA Mini Kit (QIAGEN), RNeasy Mini Kit (QIAGEN) and MagMAX™ Viral/Pathogen II Nucleic Acid Isolation Kit (Thermo Fisher Scientific). We quantified viral RNA by real time RT-PCR analysis with a one-step RT-qPCR kit (THUNDERBIRD Probe One-step RT-qPCR kit, TOYOBO) using 5’-ACAGGTACGTTAATAGTTAATAGCGT-3’ for forward primer and 5’-ATATTGCAGCAGTACGCACACA-3’ for reverse primer, and a 5’-FAM-ACACTAGCCATCCTTACTGCGCTTCG-3’ probe, as described (Corman et al., 2020). The detection limit of SARS-CoV-2 RNA in this study was 39 cycle ($C_t$ value).

*Cell viability.*

Cell viability was examined by MTT assay as previously reported (Ohashi et al., 2018) (Fig. 2C) or by quantification of survival cell numbers fixed with 4% paraformaldehyde and stained with 0.02% DAPI with a high content imaging analyzer ImageXpress Micro Confocal (MOLECULAR DEVICES) (Fig. 1C and 4A, right).

*Time-of-addition analysis.*

VeroE6/TMPRSS2 cells were inoculated with the virus at an MOI of 0.001 for 1 h, and the free virus was removed by washing. Compounds were added at three different times before measuring extracellular viral RNA (Fig. 3B): (a) throughout the entire assay covering viral lifecycle (whole: 1 h + 24 h after virus inoculation), (b) only the early phase of the assay covering viral entry steps (entry: initial 1 h + 2 h after virus inoculation), (c) during the late phase of the assay, from viral replication to virus secretion (post-entry: last 22 h after virus inoculation).

*Virus-cell attachment assay.*

Virus and compounds were preincubated at 4°C for 1 h and then exposed to VeroE6/TMPRSS2 cells at 4°C for 5 min to allow virus-cell attachment. After removing unbound virus by washing, attached viral RNA was extracted with RNeasy Mini Kit (QIAGEN) and measured by real time RT-qPCR. The same assay was done without cells to measure the background level. Heparin was used as a positive control that inhibits SARS-CoV-2 attachment to cells. The specific cell-attached SARS-CoV2 RNA was calculated by subtracting viral RNA levels without cells from those with VeroE6/TMPRSS2 cells and are shown in Fig. 3C.

*Post-attachment assay.*

The assay examining the steps from Spike cleavage/membrane fusion through viral secretion, was conducted by initiating compound treatment after viral attachment to already highly infected cells and detecting the secreted virus for a short duration (Fig. 3D): Cells were incubated with the virus at an MOI of 1.5 at 4°C for 1 h, then removing the unattached virus. These virus-attached cells were incubated at 37°C for 6 h in the presence of compounds to allow viral entry through replication and secretion. The culture supernatant was recovered to detect extracellular viral RNA. E-64d (12.5 μM), lysosomal/cytosolic cysteine protease inhibitor, was used as a positive control.

*Pseudovirus infection assay.*

SARS-CoV-2 pseudotype virus was produced using the vesicular stomatitis virus (VSV)-pseudotype system essentially as described previously (Fukushi et al., 2005; Tani et al., 2010) using the expression plasmid encoding the SARS-CoV-2 Spike protein and G-deficient VSV, which contains the luciferase gene instead of the VSV-G gene. HCV pseudotype virus was prepared from the retrovirus pseudoparticle system using the expression plasmid for murine leukemia virus Gag-Pol, luciferase protein and HCV E1E2 envelope protein (kindly provided by Dr. Francois-Loic Cosset at University of Lyon) as described (Bartosch et al., 2003).

The pseudovirus for SARS-CoV-2 was inoculated to VeroE6/TMPRSS2 cells in the presence or absence of compounds and the intracellular luciferase activity was measured at 24 h post-inoculation. Camostat (TCI) 50 μM and E-64d (Cayman) 50 μM were used as a positive control to inhibit SARS-CoV-2 entry (Fig. 3E, left). HCV pseudotype virus was inoculated to Huh-7 cells (kindly provided by Dr. Francis Chisari at The Scripps Research Institute) for 4 h, followed by washing, culturing for 72 h and measuring luciferase activity. Compounds were treated for 1 h prior to infection and for 4 h during the virus inoculation. Bafilomycin A1 at 10 ng/mL was used as a positive control for inhibiting HCV entry (Fig. 3E, right).

*Statistical analysis.*

Statistical significance was analyzed using the two-tailed Student's *t*-test (*p < 0.05; **p < 0.01; N.S., not significant).

### Supplementary Notes

*Quantification of the drug dose-response curves.*

The typical dose-response curves of a single antiviral drug can be analyzed using the following Hill function (Koizumi et al., 2017) (Fig. 2B):

$$f_u = \frac{1}{1+\left(\frac{D}{IC_{50}}\right)^m}. \quad (1)$$

Here, $f_u$ represents the fraction of infection events unaffected by the drug (i.e., $1-f_u$ equals the fraction of drug-affected events). $D$ is the drug concentration, $IC_{50}$ is the drug concentration that achieves 50% inhibition of activity, and $m$ is the slope of the dose-response curve (i.e., Hill coefficient) (Koizumi et al., 2017). Dose-response curves for drugs with higher $m$ values show stronger antiviral activity at the same normalized drug concentration when the drug concentration is higher than the $IC_{50}$ (Fig. 2B). Least-square regression approach was used to fit Eq. (1) to dose-response data and estimate the values of $IC_{50}$ and $m$. Those estimated values for each drug against SARS-CoV-2 are summarized in Table S1.

*Expected anti-SARS-CoV-2 effect of double-drug combinations by Bliss independence.*

We evaluated the effect of double-drug combinations for Bliss independence, widely used to analyze drug combination data (Bliss and Fisher, 1953; Kobayashi et al., 2014; Koizumi and Iwami, 2014; Tallarida, 2001). Bliss independence assumes that each drug acts on different targets, and is defined as:

$$f_u^{\mathrm{Bcom}} = f_u^A(D) \times f_u^B(D), \quad (2)$$

where $f_u^{\mathrm{Bcom}}$, $f_u^A$ and $f_u^B$ are the fractions of infection events unaffected by the combined drugs A (i.e., Nelfinavir: NFV) and B (i.e., Mefloquine: MFQ) expected by the Bliss model, single drug A and single drug B defined by Eq. (1), respectively. Using Eq. (2), we expected the anti-SARS-CoV2 effects of combined drugs A and B, $1-f_u^{\mathrm{Bcom}}$, from the anti-SARS-CoV-2 effects of the single drugs

*Determination of synergism.*

To determine the synergy between NEV and MFQ, both $f_u^{\mathrm{Bcom}}$ in Eq. (2) and the mean value of NEV and MFQ combination treatment data (e.g., $f_u^{\mathrm{EXcom}}$) were used to calculate $f_u^{\mathrm{Bcom}}/f_u^{\mathrm{Excom}}$. If this value is equal to 1 (i.e., $f_u^{\mathrm{Bcom}}/f_u^{\mathrm{Excom}} = 1$), then synergy is not observed (the white region in Fig. 4B). If $f_u^{\mathrm{Bcom}}/f_u^{\mathrm{Excom}} > 1$, then $0 < f_u^{\mathrm{Excom}} < f_u^{\mathrm{Bcom}} < 1$, indicating stronger antiviral activity in double combination treatment than expected by Bliss model, suggesting synergy (the orange region in Fig. 4B).

*Prediction of the antiviral effect of MFQ in clinical settings.*

We used the following simple mathematical model which was also employed in (Ohashi et al., 2020):

$$\frac{df(t)}{dt} = -\big(1 - \eta(t)\times H(t)\big)\beta f(t)V(t), \qquad (1)$$

$$\frac{dV(t)}{dt} = \big(1 - \eta(t)\times H(t)\big)\gamma f(t)V(t) - \delta V(t), \qquad (2)$$

where $f(t)$ and $V(t)$ are the ratio of uninfected target cells and the amount of virus, respectively. The parameters $\beta, \gamma$, and $\delta$ represent the rate constant for virus infection, the maximum rate constant for viral replication and the death rate of infected cells, respectively. $H(t)$ is a Heaviside step function defined as $H(t) = 0$ if $t < T$: otherwise $H(t) = 1$, where $T$ is the initiation timing of the treatment, and the anti-SARS-CoV2 effect of MFQ for $t > T$ are described as follows:

$$\eta(t) = 1 - f_u\big(D(t)\big) = 1 - \frac{1}{1 + \left(\frac{D(t)}{IC_{50}}\right)^m}, \qquad (3)$$

$$D(t) = C_{max}e^{-kt}, \qquad (4)$$

where $C_{max}$ and $k$ are the peak drug concentration and the elimination rate for the corresponding drug, respectively. We used the same values of $\beta, \gamma, \delta$, and $V(0)$ as previously defined (Ohashi et al., 2020), and the values of parameters in Eq. (3) and (4) for MFQ are summarized in Table S1 and S2. The MFQ antiviral activity was calculated with the expected pharmacokinetics in the human lung, based on the pharmacokinetics information for human peripheral blood and its distribution to the lung (Desjardins et al., 1979; Jones et al., 1994) (see Table S2).

### Supplementary Figure

### Figure. S1

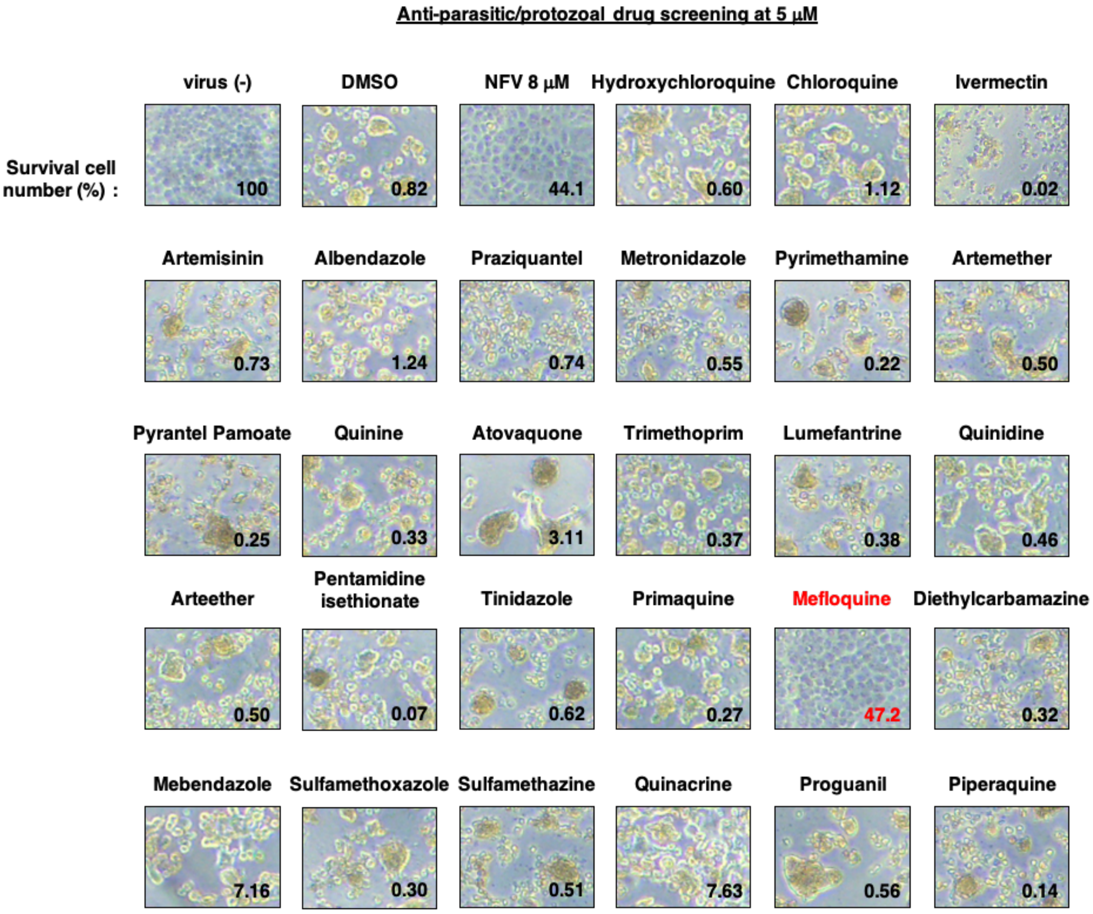


**Figure. S1.** Screening of approved anti-parasitic/anti-protozoal drugs by the cytopathic effect assay of SARS-CoV-2. VeroE6/TMPRSS2 cells were treated with 5 mM of the indicated compounds, as shown in Fig. 1A. The figures show the cells were observed by microscopy. The cell numbers were quantified with a high content imaging analyzer and are shown as the percentage of the uninfected sample [virus (-)] in the bottom of each image. NFV, suggested to inhibit SARS-CoV-2 infection, was used as positive control.

### Supplementary Tables

*Table S1. Estimated characteristic parameters of the tested antiviral drugs*

| Drug (unit) | Class | $IC_{50}$ | $m$ |
|---|---|---|---|
| Single-drug treatment | | | |
| Hydroxychloroquine ($\mu$M) | EI | 1.937 | 1.555 |
| Primaquine ($\mu$M) | unknown | 2.473 | 0.571 |
| Mefloquine ($\mu$M) | EI | 1.285 | 3.738 |
| Quinine ($\mu$M) | unknown | 14.94 | 0.573 |
| Quinidine ($\mu$M) | unknown | 3.676 | 0.564 |
| Combination treatment | | | |
| Nelfinavir ($\mu$M) | RI | 2.323 | 15.56 |
| Mefloquine ($\mu$M) | EI | 0.907 | 2.383 |

RI, replication inhibitor; EI, entry inhibitor

$IC_{50}$, 50% inhibitory concentration

$m$, the slope of the dose-response curve (i.e., Hill coefficient)

*Table S2. Summary of pharmacokinetic parameters of MFQ 1,000mg dose*

| Parameter name | Symbol | Unit | Value* |
|---|---|---|---|
| Maximum concentration | $C_{max}$ | μM | 21.6 |
| Degradation rate | $k$ | day$^{-1}$ | 0.0395 |

*Expected pharmacokinetics information in a human lung. We estimated the scaling parameter of $C_{max}$ between the lung and peripheral blood from the distribution of MFQ in the lung (Jones et al., 1994). We then calculated $C_{max}$ in the human lung multiplying that in the human peripheral blood by the scaling parameter. Since the half-life of MFQ in the lung is not available, we assumed a degradation rate of MFQ in the lung of the same value reported in the plasma (Desjardins et al., 1979; Karbwang and White, 1990).
